# Endothelial Rho kinase controls blood vessel integrity and angiogenesis

**DOI:** 10.1101/2024.11.19.624343

**Authors:** Martin Lange, Caitlin Francis, Jessica Furtado, Young-Bum Kim, James K Liao, Anne Eichmann

## Abstract

**Background:** The Rho kinases 1 and 2 (ROCK1/2) are serine-threonine specific protein kinases that control actin cytoskeleton dynamics. They are expressed in all cells throughout the body, including cardiomyocytes, smooth muscle cells and endothelial cells, and intimately involved in cardiovascular health and disease. Pharmacological ROCK inhibition is beneficial in mouse models of hypertension, atherosclerosis, and neointimal thickening that display overactivated ROCK. However, the consequences of endothelial ROCK signaling deficiency in vivo remain unknown. To address this issue, we analyzed endothelial cell (EC) specific *ROCK1* and *2* deletions.

**Methods:** We generated Cdh5-CreERT2 driven, tamoxifen inducible loss of function alleles of *ROCK1* and *ROCK2* and analyzed mouse survival and vascular defects through cellular, biochemical, and molecular biology approaches.

**Results:** We observed that postnatal or adult loss of endothelial *ROCK1* and *2* was lethal within a week. Mice succumbed to multi-organ hemorrhage that occurred because of loss of vascular integrity. ECs displayed deficient cytoskeletal actin polymerization that prevented focal adhesion formation and disrupted junctional integrity. Retinal sprouting angiogenesis was also perturbed, as sprouting vessels exhibited lack of polymerized actin and defective lumen formation. In a three-dimensional endothelial sprouting assay, combined knockdown of *ROCK1/2* or knockdown or *ROCK2* but not *ROCK1* led to reduced sprouting, lumenization and cell polarization defects caused by defective actin and altered VE-cadherin dynamics. The isoform specific role of endothelial ROCK2 correlated with ROCK2 substrate specificity for FAK and LIMK. By analyzing single and three allele mutants we show that one intact allele of *ROCK2* is sufficient to maintain vascular integrity *in vivo*.

**Conclusion:** Endothelial ROCK1 and 2 maintain junctional integrity and ensure proper angiogenesis and lumen formation. The presence of one allele of *ROCK2* is sufficient to maintain vascular growth and integrity. These data indicate the need of careful consideration for the use of ROCK inhibitors in disease settings.

## Introduction

Rho-kinase (ROCK) is a key regulator of cytoskeletal actomyosin rearrangements and contractility. Rock was identified as a serine/threonine kinase that functions downstream of the small GTPase Rho [1,2]. The two isoforms, ROCK1 and ROCK2 share 65 % overall identity in their amino acid sequence [3–5]. Their kinase domain shares 90 % amino acid identity, resulting in similar kinase activity *in vitro*. The kinase domain is followed by a coiled-coil region containing the Rho-binding domain (RBD) and a pleckstrin homology domain with a cysteine-rich domain toward the C terminus (PH-C1). Structural differences within the PH-C1 domain of ROCK1 and ROCK2 result in a binding preference to membrane lipids for ROCK2 but not for ROCK1 [6]. This leads to distinct subcellular distributions of ROCK1 and 2 that reportedly differ between various cell types [7,8].

The best-known substrate of ROCK1/2 is the myosin binding subunit (MYPT1) of the myosin light chain phosphatase (MLCP). ROCK- mediated phosphorylation of MYPT1 on Thr697 and Thr855 inhibits the catalytic activity of MLCP, leading to an increase in myosin light chain (MLC) phosphorylation [9,10]. MLC phosphorylation is responsible for cell contraction and for sustained smooth muscle contraction through calcium sensitization. Further substrates of ROCK1/2 include focal adhesion kinase (FAK) and p21-activated kinase (PAK), which regulate focal adhesions between cells and the extracellular matrix, as well as LIM-kinase (LIMK) and Cofilin that regulate actin polymerization status [11,12]. ROCK1/2 interaction with its substrates regulates cellular motility, polarity, morphology and overall gene expression [13].

Increased ROCK activity was reported in numerous cardiovascular diseases, including arteriosclerosis, restenosis, ischemia/reperfusion injury, cerebral cavernous malformation as well as cardiac hypertrophy and diastolic heart failure [14–17]. Preventing ROCK-mediated vascular hypercontractility in animal models for these diseases using pharmacological ROCK inhibitors, was shown to have beneficial effects on disease outcomes. Interestingly, the only vascular therapy applied in humans is the ROCK inhibitor Fasudil to treat cerebral vasospasm, which is approved in Japan and China [18,19]. In this disease setting, the ROCK inhibitor prevents vasoconstriction by decreasing the phosphorylation of MLC in smooth muscle cells (SMC) [20]. In the United States, the only FDA approved ROCK inhibitor application are eye drops for the treatment of glaucoma, a blinding eye disease caused by increased intraocular pressure [21]. The ROCK inhibitor helps to release intraocular pressure by acting on the trabecular meshwork and Schlemm’s canal to increase fluid drainage [22,23]. Despite these examples, ROCK inhibitor treatments have largely failed to transition to clinical applications.

To develop ROCK-inhibitor based therapies, better understanding of ROCK function in different cell types, and the role of ROCK1 and 2 isoforms in these cells *in vivo* is a major goal. *Rock2* knockout mice were described in 2003 and resulted in about 90 % lethality *in utero* [24]. Global *Rock1* knockout also resulted in 60 % lethality in utero or neonatally, depending on the genetic background. The lethality of the global knockouts was associated with defects in embryo-placenta interaction, and lack of closure of the ventral body wall [25–28]. As a result of the early lethality, studies of ROCK1 and ROCK2 in animal models of cardiovascular disease were conducted in haploinsufficient mouse models or using pharmacological ROCK inhibitors [29–33], thereby hindering understanding of cell type specificity and isoform specificity. Mouse models carrying floxed alleles of *Rock1* and *Rock2* that allow for inducible and cell-type specific deletions of these genes were generated more recently [34,35], and allowed evaluation of the role of ROCK in SMC contraction and in lymphatic endothelial cells (LECs) [8,36,37]. We previously reported that postnatal LEC-specific knockout of *Rock1* and *2* led to lethality, prompting the current investigation into endothelial ROCK function using pan-endothelial Cdh5-CreERT2 mice. We reveal the importance of *Rock1/2* in endothelial cells via genetic deletion and the involvement of ROCK in junctional integrity and angiogenesis.

## Methods

### Mice

All animal experiments were performed under a protocol approved by the Institutional Animal Care Use Committee of Yale University.

*Rock1^f/f^* mice [34], *Rock2^f/f^* mice [35] and Cdh5-CreERT2 mice [38] were previously described. Eight week old *Rock1^f/f^*, *Rock2^f/f^* or Rock1/2^f/f^ were intercrossed with Cdh5-CreERT2 mice to generate *Rock1^f/f^* Cdh5-CreERT2 (*Rock1^iEKO^*), *Rock2^f/f^* Cdh5-CreERT2 (*Rock2^iEKO^*) or *Rock1/2 ^f/f^* Cdh5-CreERT2 (*Rock1/2^iEKO^*), which were used for experiments. All mice were on C57BL/6J background and housed in pathogen-free animal facilities with a 12-hour light dark cycle. Mice of both sexes were used. Gene deletion was induced by intra-peritoneal Tamoxifen (TAM) injections (Sigma, T5648; 20 mg/ml in corn oil). Postnatal mice received 100 μg TAM daily for 3 days from P1 to P3 or P5 to P7. Adult mice received 2 mg TAM daily for 5 days starting at week 6. TAM injected CreERT2 negative littermates were used as controls. All *in vivo* experiments were performed on littermates.

### Reagents and antibodies

For immunostaining: IB4 (IsolectinB4, Invitrogen, # I21412), TER119 (Invitrogen, # 14- 5921-82), mouse VE-cadherin/CD144 (BD, # 550548), human VE-cadherin (Santa Cruz, # sc-9989), human VE-cadherin (R&D, # AF938), Phalloidin (Abcam, # ab176759), DAPI (BD, # 564907), pFAK Y576/Y577 (Cell Signaling, # 3281S), pMLC2 S19 (Cell Signaling, #3671S), ERG (Abcam, # ab196149), Podocalyxin (R&D, #AF1556), GM130 (BD, # 610822).

For Western blotting: ROCK1 (Invitrogen, # PA5-22262), ROCK2 (Invitrogen, # PA5- 78290), GAPDH (Cell Signaling, # 5174S), VE-cadherin (Santa Cruz, # sc-9989), pFAK Y576/Y577 (Cell Signaling, # 3281S), FAK (Cell Signaling, # 3285S), P120/Catenin delta1 (Cell Signaling, # 59854S), pPAK1/2 (Cell Signaling, #2605S), PAK1/2/3 (Cell Signaling, # 2604S), pLIMK1/2 (Cell Signaling, # 3841S), LIMK1 (Cell Signaling, # 3842S), pAKT (Cell Signaling, # 4060S), AKT (Cell Signaling, # 9272S), Claudin5 (Invitrogen, 35-2500), pMLC2 T18/S19 (Cell Signaling, # 95777S), MLC2 (Cell Signaling, # 3672S).

For Sulfo-NHS Biotin (S-N Biotin) injections: EZ-Link^TM^ NHS-Biotin was purchased from Thermo Fisher (# 21217) and visualized with Streptavidin Texas Red (Vector Laboratories, # 352588).

Appropriate fluorescently labeled secondary antibodies were used from Invitrogen (Alexa Fluor donkey anti-rabbit, Alexa Fluor donkey anti-goat, Alexa Fluor donkey anti-rat, Alexa Fluor donkey anti-mouse) or conjugated to horseradish peroxidase (anti-rabbit and anti-mouse IgG [H+L]; Vector Laboratories).

### Cell culture assays and siRNA transfection

Human umbilical vein endothelial cells (HUVECs) were obtained from Lonza Bioscience (# C2519A) and cultured in EGM2-Bullet kit medium (Lonza, CC-3156 & CC-4176). Depletion of *Rock1* and *Rock2* was achieved by transfecting 20 pmol of small interfering RNA (siRNA) against *Rock1* (Horizon Discovery Bioscience, # L-003536-00-0005) or *Rock2* (Horizon Discovery Bioscience, # L-004610-00-0005) using Lipofectamine RNAiMax (Invitrogen, # 13778150). Knockdown efficiency was assessed by Western blotting. Experiments were performed 60-72 h post-transfection and results were compared with ON-TARGETplus Non-targeting Pool (Horizon Discovery, # D-001810-10- 05).

For calcium switch assay HUVECs were treated with siRNA and grown until completely confluent. Once confluent, they were treated with EGTA (3mM final concentration) for 1 hour as described [39]. Next, cells were washed 3x with PBS, incubated in EGM-2 media and fixed at 0-, 30- and 60-minutes post treatment. VE-cadherin immunostaining was measured by microscopy; images were thresholded consistently for each group. VE-cadherin fluorescent intensity was measured for the whole cell as well as intensity at the cell membrane.

Lentivirus was generated via the 2^nd^ generation lentiviral system, using one transfer plasmid, one packaging plasmid (psPAX2, Addgene: #12260) and one envelope plasmid (pVSVG, Addgene: #8454). Plasmids were transfected in a 3:1 PEI:DNA ratio into 90% confluent HEK293 cells in a 10cm dish (PEI=polyethylenimine, Sigma 408727). On the day of transfection, the HEK293 cells were treated with 50uM Chloroquine (Sigma C6628). Media was changed at 4-hours post-transfection and cells were then left to incubate for 72 hours. After media collection, virus was concentrated and stored at -80 C [40]. HUVECs were transduced during bead coating with 10ng/mL polybrene (Fisher Scientific TR1003G). Both the LifeAct-GFP and VE-cadherin-GFP transfer plasmids were purchased from Addgene (LifeAct: 58470, VECAD: 118732).

The fibrin-bead sprouting assay was performed as reported by Nakatsu *et al.* [41]. Briefly, HUVECs were seeded onto microcarrier beads followed by siRNA-treatment and/or viral transduction. The following day, coated microbeads were embedded in a fibrin matrix. Once the clot was formed, media was overlaid along with 100,000 normal human lung fibroblasts. Sprout numbers were determined by counting the number of multicellular sprouts emanating from individual microcarrier beads. Sprout lengths were determined by measuring the length of a multicellular sprout beginning from the tip of the sprout to the microcarrier bead surface. Percent of lumenized sprouts were determined by quantifying the proportion of multicellular sprouts whose length was <80% lumenized. Sprout widths were determined by measuring the sprout width at the midpoint between the tip and the microcarrier bead. Experimental repeats are defined as an independent experiment in which multiple cultures containing sprouting beads were quantified.

### Immunostaining

For retinas: The eyes of P4/P5/P6 pups were fixed in 4% Formaldehyde for 8 min at room temperature. Retinas were dissected, blocked for 30 min at room temperature in blocking buffer (5 % non-immune donkey serum, 1 % BSA, 0.1 % Triton X-100, 0.05 % Sodium Azide in PBS at pH 7.4) and then incubated with specific antibodies in blocking buffer overnight at 4°C. The next day, retinas were washed and incubated with IB4 together with the corresponding secondary antibody overnight at 4°C, then washed and post-fixed with 0.1% PFA and mounted in DAKO fluorescent mounting medium (Agilent, # S302380-2). High-resolution pictures were acquired using ZEISS LSM-780, Leica SP8 inverted or an inverted Nikon Eclipse Ti2 microscope. Quantification of vascular outgrowth as well as was quantification of pixel intensity were performed using the software Image J. Vascular density was quantified with the software Angiotool by quantifying the vascular surface area normalized to the total surface area. Quantification of pixel intensity and image processing was performed using the software ImageJ and Imaris.

For HUVECs: Cells were plated on gelatin coated dishes. Confluent cells were fixed for 10 min with 4% Formaldehyde and permeabilized with 0.1% Triton X- 100 for 10 min followed by blocking for 30 min in blocking buffer (5 % non-immune donkey serum, 1 % BSA, 0.1 % Triton X-100, 0.05 % Sodium Azide in PBS at pH 7.4). Afterwards, cells are incubated with primary antibody for 1 hour and then secondary antibody conjugated with fluorophore for 1 hour. Finally, cells are mounted in DAKO fluorescent mounting medium (Agilent, # S302380-2).

For aorta *en face*: Mice were anesthetized and perfused with PBS followed by perfusion with 4 % Formaldehyde. Afterwards aorta was harvested and fixed in 4 % Formaldehyde. After removing fat tissue and connective tissue the aorta is cut longitudinally and pinned down with insect needles. Next, aortas are blocked for 1 hour at room temperature in blocking buffer (5 % non-immune donkey serum, 1 % BSA, 0.1 % Triton X-100, 0.05 % Sodium Azide in PBS at pH 7.4) and then incubated with specific antibodies in blocking buffer overnight at 4°C. The next day, aortas were washed and incubated with corresponding secondary antibody as well as DAPI and phalloidin for 2 hours. After washing, the aortas are mounted in DAKO fluorescent mounting medium (Agilent, # S302380-2).

### Western blotting

Cells were lysed with Laemmli sample buffer (BIORAD, # 1610747) including phosphatase and protease inhibitors (Thermo Scientific, # 78420; Roche, # 11836170001). Protein lysates were separated on 4% - 15% Criterion precast gels (Biorad, # 5678084) and transferred on 0.45 µm nitrocellulose membranes (Biorad, # 1620115). Western blots were developed with Pierce ECL Western blotting substrate (Thermo Scientific, # 32106) on a Luminescent image Analyzer, Biorad ChemiDoc XRS+ imaging system. Bands were quantified using ImageJ.

### Intraperitoneal injection of SulfoNHS Biotin (S-N Biotin)

S-N Biotin tracer was injected intra-peritoneally in P5 mice at a concentration of 0.5 mg/g and left to circulate for 1 hour. Afterwards, retinas were dissected and processed as described above. Retinas were stained with IB4 and S-N Biotin perfusion was visualized using Streptavidin Texas Red (Vector Laboratories, # 352588).

### Bulk RNA-sequencing on HUVECs

Total mRNA was extracted from HUVECs using RNeasy Plus Mini Kit (QIAGEN, # 74134). RNA concentration was measured, and RNA integrity was determined using Bioanalyzer (Agilent). Using 300ng of RNA per sample, libraries were constructed using the KAPA Stranded mRNA-Seq Kit (Illumina Platforms; Kapa Biosystems), with 10 cycles of PCR amplification followed by purification using AxyPrep Mag PCR Clean-up kit (AygenTM). mRNA libraries were quantified using a Qubit fluorometer (Life Technologies) and sequenced on an Illumina® Hiseq 2500 instrument (Illumina, San Diego, CA, USA) by paired-end sequencing. Downstream analysis was performed using Partek Flow software (build version 12.2.2) (Partek, Inc, St. Louis, MI). Adapter sequences and low-quality bases were trimmed. The trimmed reads were then aligned to the hg38 reference genome using STAR and quantified to an annotation model. Gene counts were then normalized in Partek Flow. DESeq2 was then used for differential gene expression analysis to compare *siRock1* + *siRock2* samples and samples transfected with control siRNA and gene expression levels were compared using a volcano plot.

### Statistical analysis

Analysis of live VE-cadherin dynamics was performed using Imaris spots tool. Spots were generated and analyzed for movement over time. Drift was corrected to avoid any quantification of unrelated signal movement. For analysis of live actin dynamics, due to a high background all analysis was performed manually. Statistical analysis was performed using GraphPad Prism 8 software. Mann-Whitney U test was used for statistical analysis of two groups. ANOVA followed by Bonferroni’s multiple comparisons test was applied for statistical analysis between 3 or more groups. Mantel-cox test was performed for survival curve statistical analysis.

## Results

### Endothelial *Rock* deletion is lethal due to in multi-organ hemorrhage

To investigate the role of endothelial ROCK *in vivo*, we generated endothelial cell-specific *Rock* double mutant mice by crossing Rock1^flox/flox^ [34] and *Rock2^flox/flox^* mice [35] with Cdh5-CreERT2 mice [38] (hereafter *Rock1/2^iEKO^*). We induced gene deletion in neonates by Tamoxifen (TAM) injection (100 μg daily) at postnatal day (P) 5-P7 and monitored mouse survival (Figure 1 A). All 8 *Rock1/2^iEKO^*mutant mice died around P12, while all 8 TAM injected littermate controls survived (Figure 1 B). Likewise, when gene deletion was induced in adult 8 weeks old mice, all 5 *Rock1/2^iEKO^* mice died about a week after the last TAM injection, while the 5 Cre negative control littermates survived (Figure 1 C, D). Western blot analysis of protein extracts derived from purified mouse brain ECs of P4 mice revealed efficient EC deletion of *Rock1/2* (Figure S1 A), demonstrating that endothelial ROCK1/2 are necessary for mouse survival. To determine the cause of death, *Rock1/2^iEKO^* mice injected at P5-P7 were sacrificed at P10 and internal organs were inspected. We observed multiple hemorrhages in the brains and retinas of *Rock1/2^iEKO^* mice but not in littermate controls (Figure 1 E and F). Hemorrhages in *Rock1/2^iEKO^* mice but not in littermate controls were also visible in the stomach and throughout the gastrointestinal (GI) tract including the small intestine, caecum, and large intestine (Figure 1 G). In addition, cryosections of tissue from the kidney, heart and liver were stained with IsolectinB4 (IB4) to visualize blood vessels and TER119 to label erythrocytes. In the control mice, TER119 positive erythrocytes could only be seen within capillaries and larger vessels, while erythrocytes in *Rock1/2^iEKO^* mice were observed to accumulate outside of blood vessels in all three organs (Figure1 H). These results show that *Rock1/2^iEKO^* mice succumb to multi organ hemorrhages.

**Figure 1:**
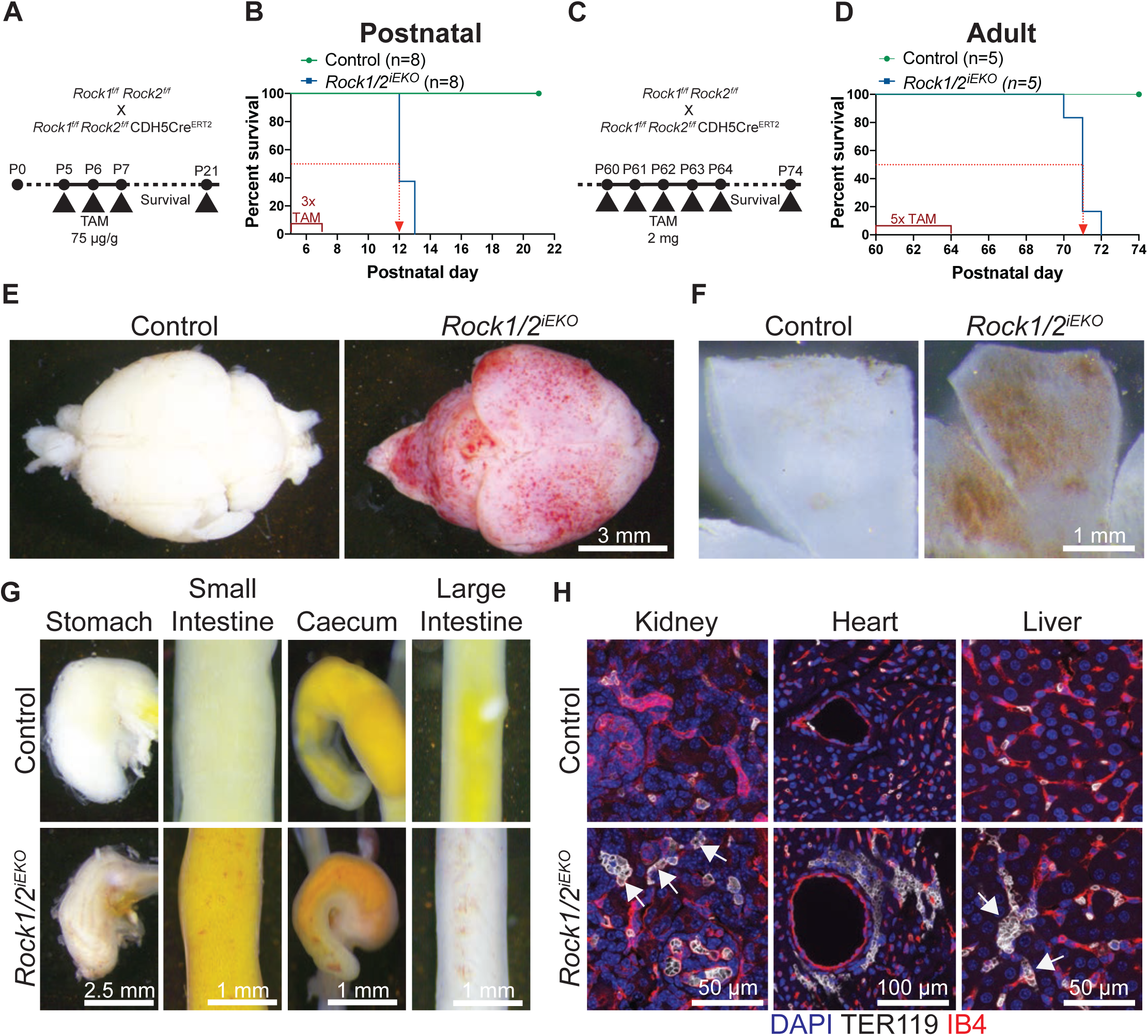
Endothelial *Rock1/2* deletion is lethal due to multi-organ hemorrhage. (A) *Rock1/2* gene deletion strategy using TAM injection in postnatal mice. (B) Survival curve for postnatal *Rock1/2^iEKO^* mice. (C) *Rock1/2* gene deletion strategy using TAM injection in adult mice. (D) Survival curve for adult *Rock1/2^iEKO^* mice; green line represents control mice (Cre negative littermates); blue line presents mutant mice, and red dotted line indicates timepoint of 50 % survival. (E-F) Brightfield images of mouse brains (E) and retinas (F) of control mice and *Rock1/2^iEKO^* mice at P10. (G) Brightfield images of mouse GI tract at P10. (H) Immunofluorescence staining with IB4 (red – blood vessel ECs), TER119 (white – erythrocytes) and DAPI (blue – nuclei) of mouse tissues from kidney, heart and liver at P10; comparing control and *Rock1/2*^iEKO^.

### Defective actin polymerization and junction defects in *Rock1/2^iEKO^* mice

To investigate how actin is affected in ECs of *Rock1/2^iEKO^* mice, we used aorta *en face* preparations from 8 week old mice. Aortic ECs were stained with phalloidin to label F-actin and an antibody recognizing VE-cadherin. In the controls, phalloidin robustly labeled the cortical actin as well as stress fibers, and VE-cadherin was evenly distributed along the cell junctions (Figure 2 A). By contrast in the *Rock1/2^iEKO^* mice, the phalloidin staining was much weaker and the VE-cadherin positive cell junctions were irregular in appearance and showed numerous gaps between cells (Figure 2 A). Transfection of human umbilical vein endothelial cells (HUVECs) with control or *ROCK1/2* siRNA produced a similar phenotype, in that control siRNA transfected HUVECs showed strong phalloidin staining and evenly distributed VE-cadherin at cell-cell junctions, while *ROCK1/2* knockdown HUVECs showed weaker phalloidin staining and disturbed irregular VE-cadherin labeled cell-cell junctions with gaps between some of the cells (Figure 2 B). Western blotting showed efficient knock down of *ROCK1* and *ROCK2* normalized to the housekeeping gene GAPDH and a clear reduction of pMLC (Figure 2 C-F). Additional Western blot analysis of ROCK-dependent cytoskeletal regulatory proteins LIMK and Cofilin revealed reduced phosphorylation in *ROCK1/2* knockdown cells compared to controls (Figure 2 C, G-H), along with decreased phosphorylation of Focal adhesion kinase (FAK) and PAK1/2 (Figure 2 C, I-J). Staining of HUVECs with pFAK antibody and phalloidin further revealed decreased amounts of polymerized actin and few remaining focal adhesions at the tips of actin strands (Figure 2 O). Interestingly, there was no significant difference between the overall amounts of VE-cadherin or P120 in control and *ROCK1/2* knockdown HUVECs (Figure 2 C, K-L), despite the observed defects in adherens junctions in the *ROCK1/2* deficient endothelium. However, the phosphorylation of AKT and the overall amounts of Claudin-5 were decreased (Figure 2 C, M-N).

**Figure 2:**
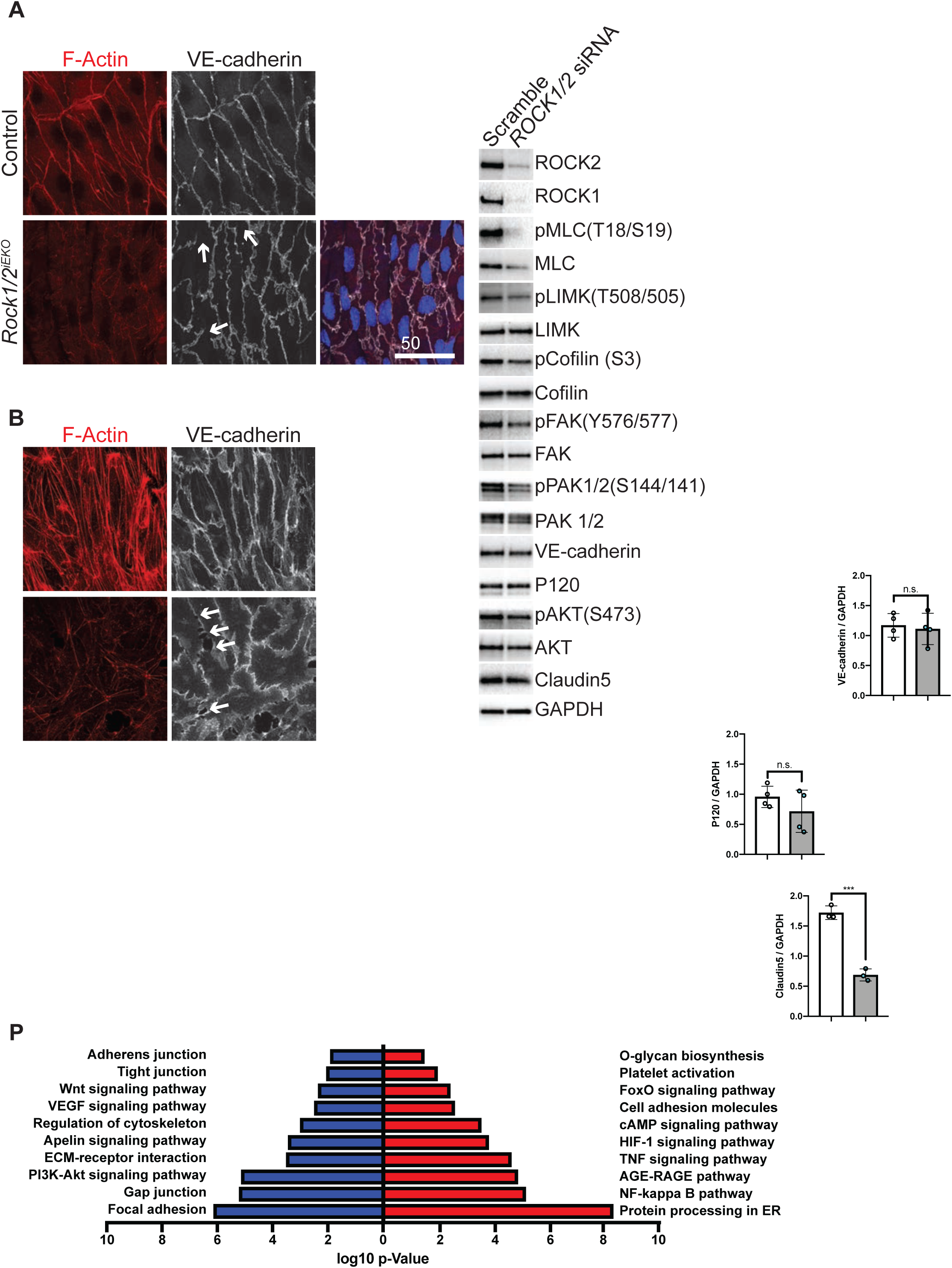
Defective actin polymerization and junctional defects in *Rock1/2^iEKO^* mice. (A-B) Staining with Phalloidin, VE-cadherin and DAPI of adult mouse aorta after *en face* preparation (A) and HUVECs after *ROCK1/2* siRNA or control siRNA treatment (B); white arrows point to disrupted junctions. (C-N) Western blot analysis (C) and quantification (D-N) of HUVEC protein extracts. Quantification of ROCK1 (D), ROCK2 (E), pMLC (F), pLIMK (G), pCofilin (H), pFAK (I), pPAK (J), VE-cadherin (K), P120 (L), pAKT (M) and Claudin5 (N) was performed by normalizing values to GAPDH or total form of respective kinase for control siRNA treated HUVEC and *ROCK1/2* siRNA treated HUVEC. Dots represent individual experiments; black lines indicate the mean value s.d.; n.s = not significant; *p<0.05; **p<0.01; ***p<0.001. Groups were compared by Mann-Whitney U test. (O) HUVECs after *ROCK1/2* siRNA or control siRNA treatment stained with Phalloidin, pFAK and DAPI. (P) Pathway enrichment analysis of up- and down-regulated genes between control siRNA treated HUVEC and *ROCK1/2* siRNA treated HUVEC (n=3/group) based on log2 fold-change against -log10 (*p*-value). Red color indicates activation and blue color indicates suppression.

As knockdown in HUVECs phenocopied cytoskeletal and junctional defects observed in *Rock1/2^iEKO^* mice, we isolated mRNA for RNA sequencing and compared control siRNA treated HUVECs with *ROCK1/2* siRNA treated HUVECs. Down-regulated pathways included adherens junctions, tight junctions, gap junctions and focal adhesions, as well as actin cytoskeleton regulation, growth factor signaling and ECM-receptor interaction (Figure 2 P, Figure S1 B). Pathways associated with up-regulated genes included NF-kappa B signaling, HIF-1 signaling pathway as well as cell adhesion molecules (Figure 2 P). Hence, endothelial *ROCK* deletion decreased transcription of pathways that control endothelial barrier function, which may further amplify loss of polymerized actin and focal adhesions and thereby trigger endothelial barrier breakdown and hemorrhage.

### ROCK promotes retinal blood vessel integrity

To understand how the hemorrhagic phenotype developed within organs, we performed a time-course of *Rock1/2* deletion and analyzed postnatal retina development. The entire vasculature of this organ can be visualized in a single whole-mount preparation, allowing us to detect when and where changes in vascular integrity occur. *Rock1/2^iEKO^* mice were injected with 100 µg TAM at P1, P2 and P3 and analyzed at P4, P5 and P6 (Figure 3 A-D). The retinas were stained with IB4 to label blood vessels and TER119 antibodies to label erythrocytes. Hemorrhages were quantified by measuring the area occupied by extravascular erythrocytes and expressed as percentage of the total vascular IB4+ area. At P4, no differences in extravasated erythrocytes were observed between *Rock1/2^iEKO^* and control littermates and the few extravasated erythrocytes were located at the leading edge of the developing retina in both genotypes (Figure 3 A-B), which is known to be leaky [42]. As early as P5, hemorrhagic foci formed at the vascular front in *Rock1/2^iEKO^* mice, which further increased in severity at P6 (Figure 3 A, C and D). Staining the retinas with IB4 and either phalloidin or pMLC confirmed that the retinal vasculature of *Rock1/2^iEKO^* mice showed decreased phalloidin and pMLC staining compared to control mice (Figure 3 E and F). VE-cadherin staining revealed disturbed junctions and changes in VE-cadherin distribution along the cell junctions in *Rock1/2^iEKO^*retina vessels (Figure 3 G). In addition, angiogenic sprouts at the vascular front of *Rock1/2^iEKO^* mice showed a significant reduction of pFAK staining compared to littermate controls (Figure 3 H).

**Figure 3:**
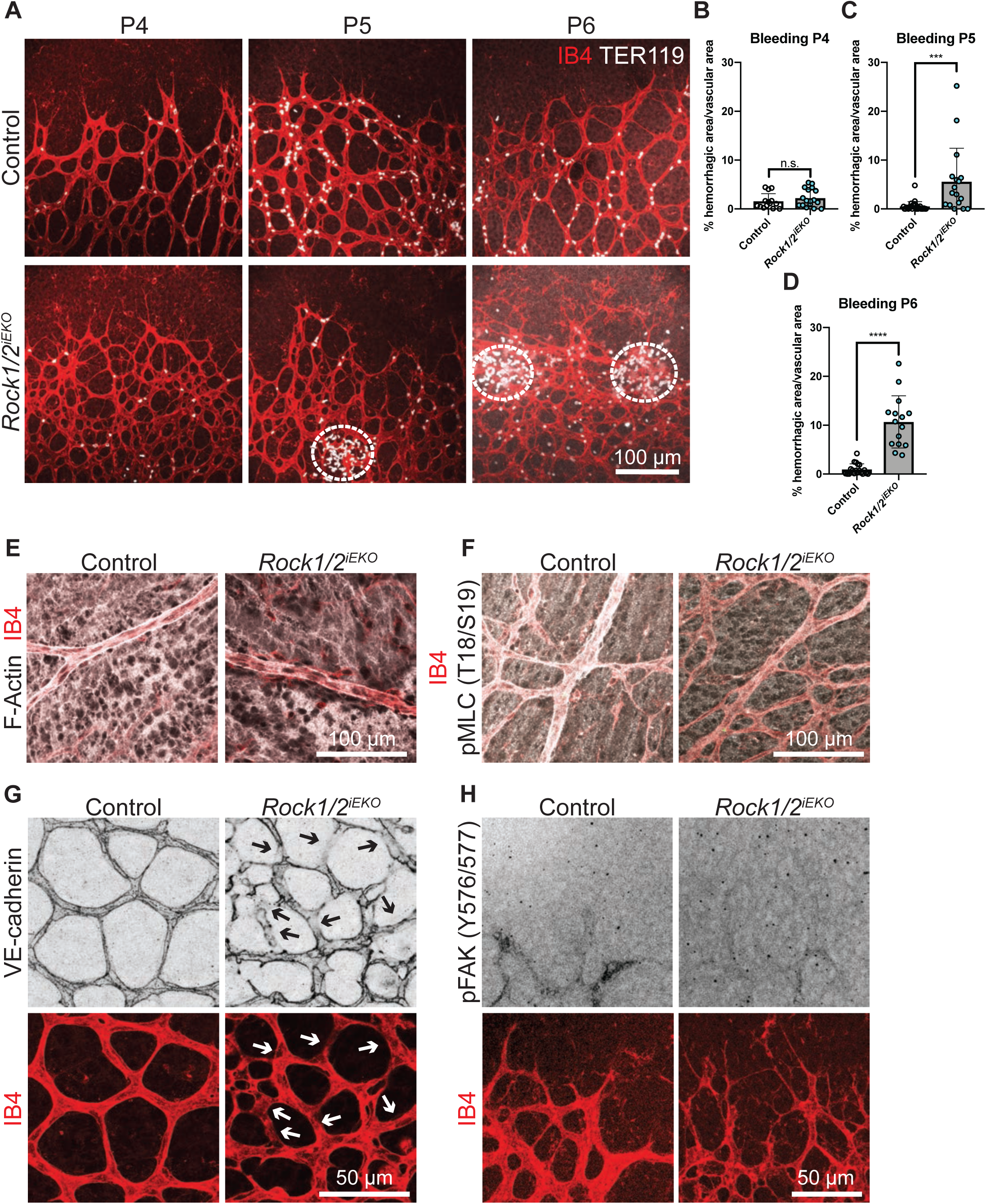
ROCK promotes retinal blood vessel integrity. (A) Whole mount retinas from control and *Rock1/2^iEKO^* mice at P4, P5 and P6, stained with IB4 and TER119; stippled circles indicate extravascular erythrocytes. (B-D) Quantification of bleedings presented in % of extravascular TER119, at P4 (B), at P5 (C) and P6 (D) for control and *Rock1/2*^iEKO^. Dots represent individual retinas; black lines indicate the mean value s.d. n.s = not significant; ***p<0.001; ****p<0.0001. Groups were compared by Mann-Whitney U test. (E) P5 retinas of control and *Rock1/2^iEKO^* mice, stained with IB4 and Phalloidin. (F) Whole mount retinas from control and *Rock1/2^iEKO^* mice at P5, stained with IB4 and pMLC. (G) Whole mount retinas from control and *Rock1/2^iEKO^* mice at P5, stained with IB4 and VE-cadherin; Arrows point to missing or disrupted junctions. (H) Whole mount retinas from control and *Rock1/2^iEKO^* mice at P5, stained with IB4 and pFAK.

### ROCK controls retinal angiogenesis and lumen formation

In addition to hemorrhage, we also noted angiogenic defects in *Rock1/2^iEKO^* mice. Compared to control littermates, vascular outgrowth in *Rock1/2^iEKO^* retinas was reduced by 20 % (20.43 ± 1.5) at P6 (Figure 4 A-B). To understand how ROCK1/2 affects retina angiogenesis, we co-stained vessels with IB4 and ERG to quantify endothelial nuclei within the growing blood vessels. Compared to the retinas of control mice, the *Rock1/2^iEKO^* mice displayed reduced numbers of EC nuclei, resulting in a vascular front with few nuclei and thin blood vessels branches (Figure 4 C). Dropout of *Rock1/2* deficient endothelial nuclei started at P4 and further aggravated at P5 and P6, accompanied by an increased number of thin vascular networks (Figure 4 C-F). This leads to an overall increased IB4 positive vascular front area (Figure S2 A-B), increased branch points (Figure S2 C) and increased number of filopodial extensions (Figure S2 D). Injection with Sulfo-NHS Biotin (S-N Biotin) tracer revealed that all IB4-positive vessels in controls were perfused with S-N Biotin up to the leading tip cells (Figure 4 G). However, in the *Rock1/2^iEKO^* mutant retinas the IB4-positive vascular front was mostly unperfused (Figure 4 G). In addition, staining with the luminal marker Podocalyxin revealed areas that were positive for IB4 but negative for Podocalyxin in *Rock1/2^iEKO^* mutant retinas, while all IB4-positive areas were positive for Podocalyxin in controls (Figure 4 H). The hemorrhages in *Rock1/2^iEKO^* mutants occurred at the junctions between perfused and unperfused hyperbranched membrane extensions at the vascular front (see Figure 3A). Staining with cleaved Caspase-3 to detect apoptotic cells revealed no differences in vascular apoptosis between control and *Rock1/2^iEKO^* P5 mutant retinas (Figure S2 E-F), and ERG and KI67 staining showed that the percentage of proliferating ECs did not differ between genotypes (Figure S2 G-H). This shows that loss of ECs was not due to increased cell death or decreased proliferative capacity, but to defective sprouting angiogenesis and lumen formation at the vascular front.

**Figure 4:**
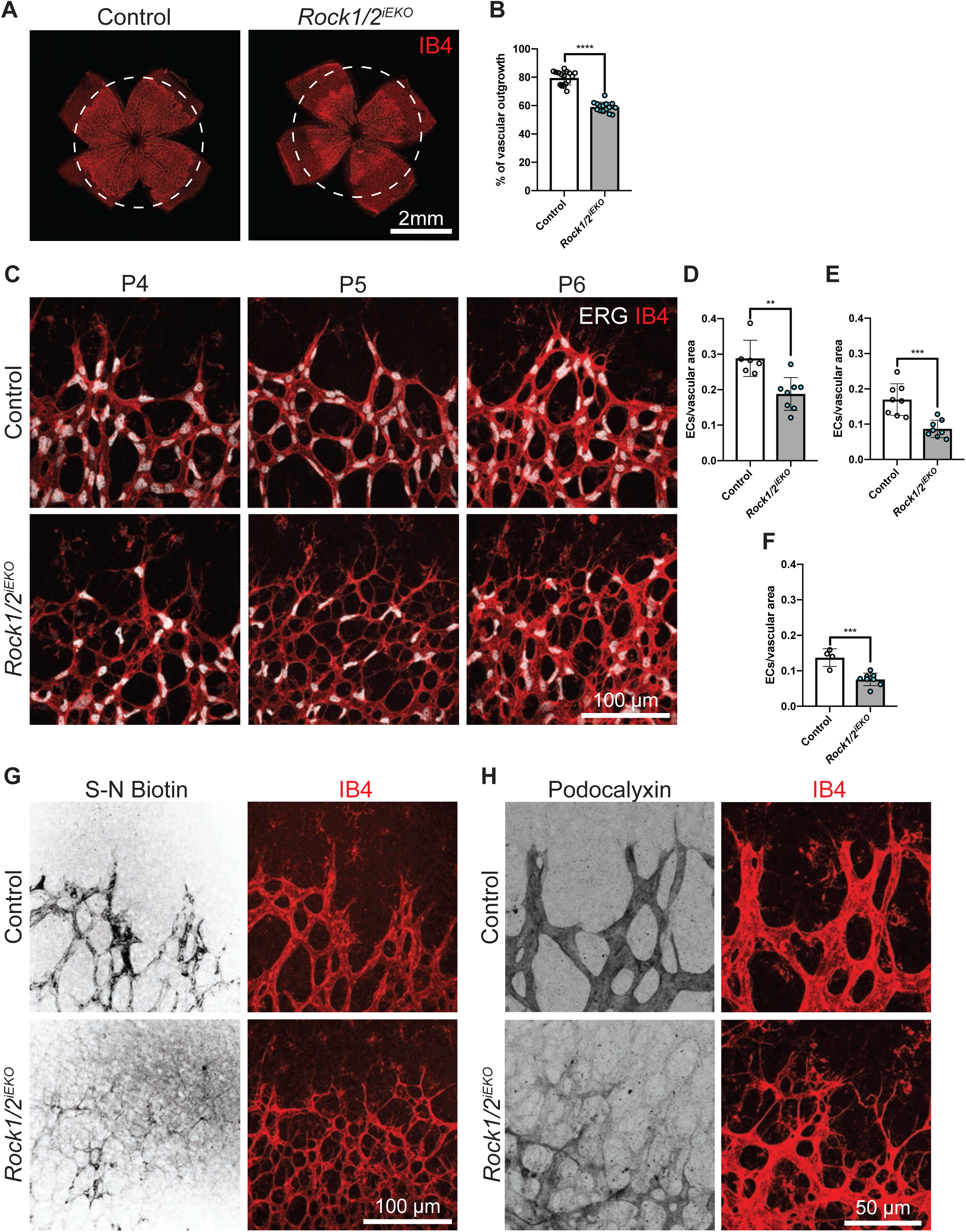
ROCK controls retinal angiogenesis and lumen formation. (A) Whole mount retinas from control and *Rock1/2^iEKO^* mice at P6 stained with IB4. (B) Quantification of vascular outgrowth at P6 for control and *Rock1/2^iEKO^*. (C) Whole mount retinas from control and *Rock1/2^iEKO^* mice at P4, P5 and P6 stained with IB4 and ERG. (D-F) Quantification of endothelial ERG+ nuclei per IB4+ vascular area, at P4 (D), P5 (E) and P6 (F) for control and *Rock1/2^iEKO^*. Black lines indicate the mean value s.d. **p<0.01; ***p<0.001; ****p<0.0001. Groups were compared by Mann-Whitney U test. (G) Whole mount retinas from control and *Rock1/2^iEKO^* mice 1 h after i.p. injection with S-N Biotin at P5 stained with IB4 and Streptavidin. (H) Whole mount retinas from control and *Rock1/2^iEKO^* mice at P5 stained with IB4 and Podocalyxin.

### *ROCK* isoform knockdown affects *in vitro* sprouting angiogenesis

To understand how ROCK promotes angiogenesis and to define how individual ROCK1 and 2 isoforms affect this process, we examined the effects of *ROCK* siRNA knockdown (KD) in HUVECs in the sprouting bead assay [41]. We targeted *ROCK1*, *ROCK2* and combination *ROCK1/2* and challenged vessels to grow by a combination of growth factors contained in the medium and secreted by fibroblast included within the beads. Loss of *ROCK1* resulted in a modest reduction of vessel length whereas KD of *ROCK2* and combination *ROCK1/2* resulted in a significant loss of sprouts per bead, vessel length, and nuclei per sprout (Figure 5 A-E). Immunohistochemical staining of VE-cadherin and F-actin showed a progressively less robust network of actin, thinner appearing junctions, and a much narrower lumen from *ROCK1* to *ROCK2* and to *ROCK1/2* knockdown (Figure 5 F-H). In all parameters measured, combination *ROCK1/2* consistently resulted in the greatest defects. These results suggest that loss of ROCK2 causes greater deficiencies relative to ROCK1, and that these deficiencies are more pronounced with a combined loss of ROCK1/2.

**Figure 5:**
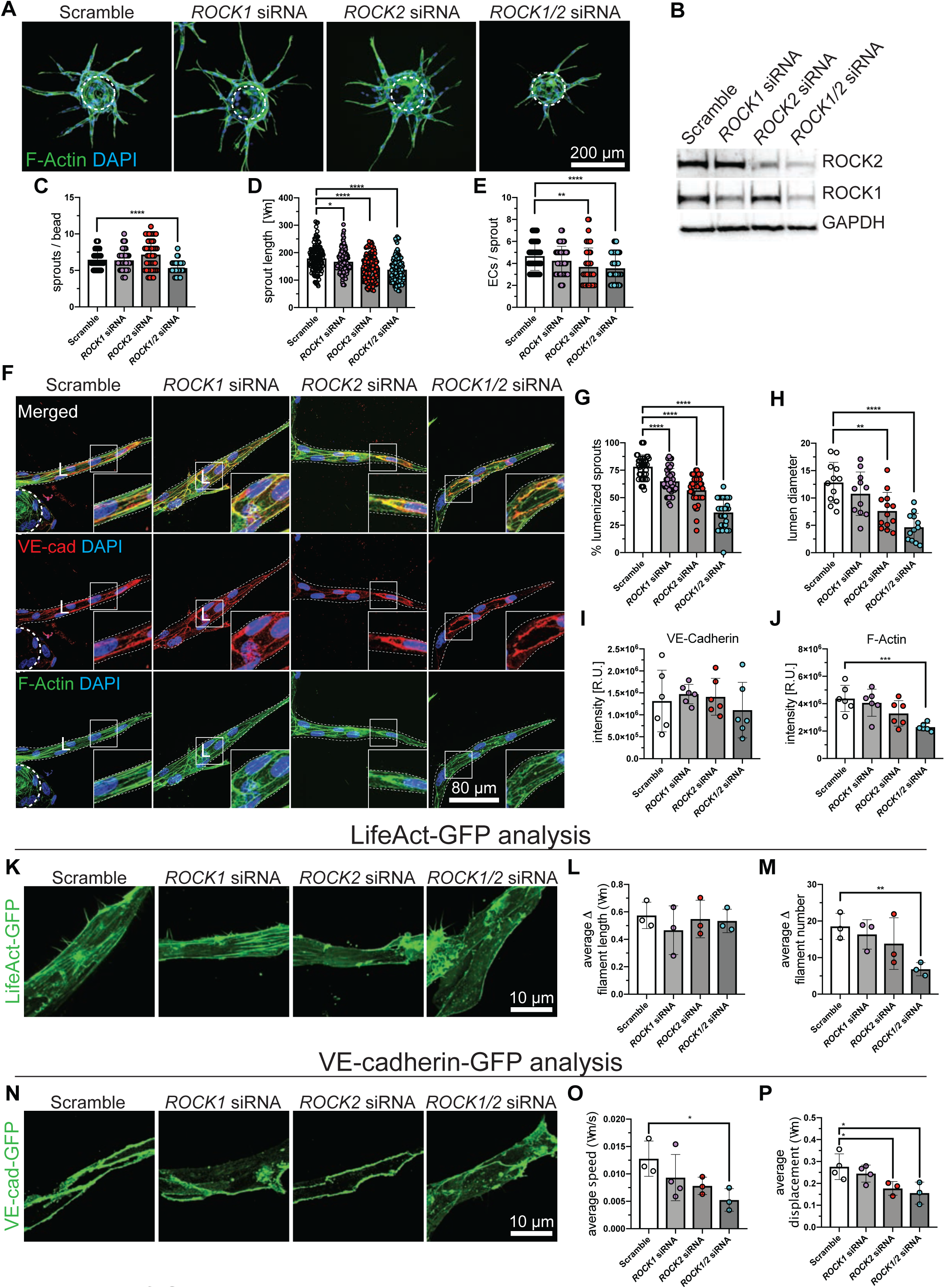
ROCK isoform knockdown affects in vitro sprouting angiogenesis. (A) Overview pictures of human endothelial vessel sprouting during fibrinogen bead assay stained with Phalloidin, and DAPI *ROCK1* and *ROCK2* knock down. Dashed circle denotes bead. (B) Western blot analysis of protein extracts from fibrinogen bead assay. (C-E) Quantification of sprouts per bead (C), sprout length (D) and nuclei per sprout (E). Dots represent individual beads; black lines indicate the mean value s.d.; *p<0.05; **p<0.01; ****p<0.0001; Groups were compared by ANOVA multiple comparisons test. (F) Closeup pictures of individual sprouts stained with Phalloidin, VE-cadherin and DAPI after *ROCK1* and *ROCK2* knock down. (G-J) Quantification of lumenized sprouts, lumen diameter, VE-cadherin fluorescent intensity and F-Actin fluorescent intensity. Dots represent individual beads or sprouts; black lines indicate the mean value s.d.; **p<0.01; ***p<0.001; ****p<0.0001; Groups were compared by ANOVA multiple comparisons test. (K) Still images of live imaging actin for different knock down conditions. (L-M) Quantification of average change in filament length over time (L) and average change in filament number over time (M). n=average of ten 1- minute interval timepoints (n= 3 sprouts for each condition); black lines indicate the mean value s.d.; **p<0.01; Groups were compared by ANOVA multiple comparisons test. (N) Still images of live VE-cadherin for different knock down conditions. (O-P) Quantification of average speed of VE-cadherin tracks over time (O) and average displacement over time. n= at least 3 sprouts for each condition; black lines indicate the mean value s.d.; *p<0.05; Groups were compared by ANOVA multiple comparisons test.

To better dissect the influence of ROCK on endothelial junctions, we measured the fluorescent intensity of endogenous F-actin and VE-cadherin in vessel sprouts. There was no significant difference in VE-cadherin. However, F-actin showed significantly less signal in *ROCK1/2* KD vessel sprouts (Figure 5 I-J). Although there was no difference in intensity of VE-cadherin, we reasoned there may be differences in junctional dynamism. Therefore, we generated GFP-tagged VE-cadherin and LifeAct lentivirus and transduced HUVECs to permit for live imaging of junctional and F-actin dynamics. Live imaging revealed less mobile F-actin and VE-cadherin in *ROCK2* and *ROCK1/2* knockdown sprouts. Like our observations in fixed sprouts, the F-actin network was less dense and VE-cadherin less robust (Figure 5 K-M). Quantification of number of filaments over time demonstrated that F-actin is indeed less filamentous in *ROCK1/2* KD vessels (Figure 5 L-M). Of note, ROCK2 F-actin was visibly less filamentous, however quantification of overall fluorescence intensity per sprout was not significantly different from the control as ROCK2 deficient actin accumulated at the plasma membrane. Analysis of live VE-cadherin demonstrated a significant reduction in speed and displacement in *ROCK1/2* vessel sprouts (Figure 5 N-P). These results suggest that loss of ROCK causes a reduction in the F-actin network and subsequent junctional instability.

### Substrate specificity of ROCK isoforms in ECs

As turnover of junctional proteins requires cooperation with the actin cytoskeleton and is imperative to vessel sprouting as well as vessel maintenance, we sought to determine if turnover of VE-cadherin was also affected with loss of *ROCK* by performing a junctional reformation assay. We treated HUVECs with EGTA, washed out the EGTA and fixed cells after 0, 30, and 60 minutes of recovery. Results of this assay indicate that junctions are slower to reform after treatment in *ROCK2* and *ROCK1/2* KD cells, while *ROCK1* KD cells behaved like control cells (Figure 6 A-I). In addition, we used a scratch wound migration assay in cells stimulated with VEGF-A (Figure 6 J-K). *ROCK1* KD cells and controls both polarized towards the leading edge, while *ROCK2* and *ROCK1/2* KD HUVECs failed to polarize towards the migration front.

**Figure 6:**
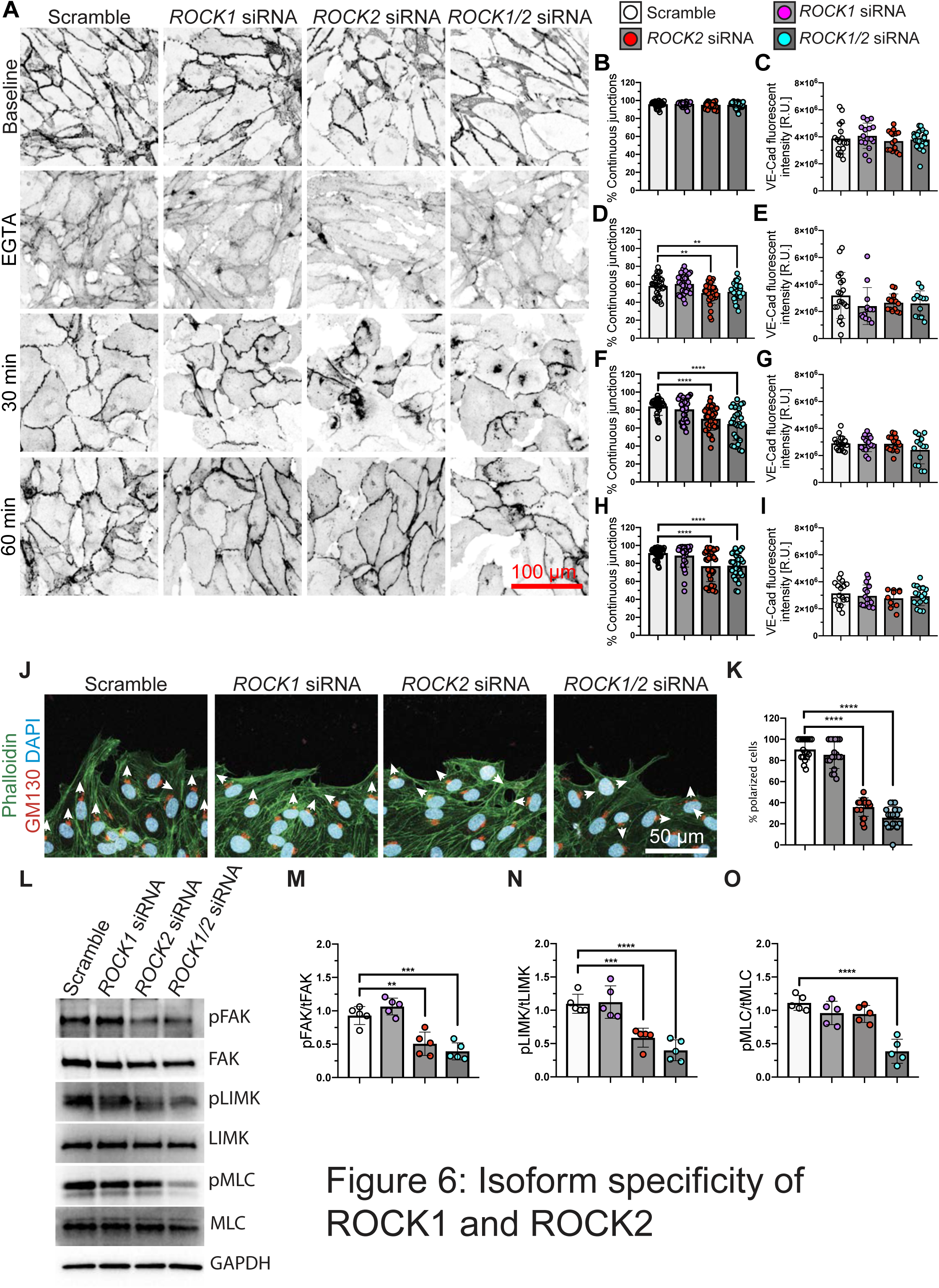
Substrate specificity of Rock isoforms in ECs. (A) HUVECs after control siRNA, *ROCK1* siRNA, *ROCK2* siRNA or *ROCK1/2* siRNA treatment stained with VE-cadherin during different conditions of the calcium switch assay. (B-I) Quantification of % continuous junctions (B, D, F and H) and VE-cadherin fluorescent intensity (C, E, G and I). (J) HUVECs after control siRNA, *ROCK1* siRNA, *ROCK2* siRNA or *ROCK1/2* siRNA treatment stained with Phalloidin, GM130 and DAPI. (K) Quantification of cell polarization after VEGFA stimulation in HUVEC with control siRNA, *ROCK1* siRNA, *ROCK2* siRNA or *ROCK1/2* siRNA treatment. Cell were counted as polarized if the Golgi is within a 90° angle to the front. Results are presented in percentage of polarized cells. Dots represent individual pictures (n = 18 for each condition); black lines indicate the mean value s.d.; ****p < 0.0001; Groups were compared by ANOVA multiple comparisons test. (L-O) Western blot (L) and quantifications (M-O) of HUVEC protein extracts after *ROCK1*, *ROCK2* and *ROCK1/2* knock down. Quantification of pFAK (M), pLIMK (N) and pMLC (O) was performed by normalizing values to total form of respective kinase for each condition. Dots represent individual experiments; black lines indicate the mean value s.d. **p<0.01; ***p < 0.001; ****p < 0.0001; Groups were compared by ANOVA multiple comparisons test.

Our in vitro results suggested that ROCK2 played a more important role compared to ROCK1, suggesting the existence of ROCK2 specific targets. To identify substrates specific to ROCK1 and 2, we used HUVECs in combination with either *ROCK1* siRNA, *ROCK2* siRNA or *ROCK1/2* siRNA to obtain protein extracts (Figure 6 L-O). Interestingly, the phosphorylation of FAK and LIMK was decreased only when *ROCK2* was knocked down, but not under *ROCK1* knock down condition (Figure 6 L and M-N). However, the phosphorylation of MLC was only affected when both *ROCK* isoforms were knocked down (Figure 6 J and O).

### Isoform specific deletion of R*OCK* in endothelial cells

To understand isoform specific roles of ROCK1 and ROCK2 *in vivo*, we generated *Rock1^flox/flox^* and *Rock2^flox/flox^* mice with Cdh5-CreERT2 mice (hereafter *Rock1^iEKO^* and *Rock2^iEKO^*). TAM injection at P5-P7 in *Rock1^iEKO^* and *Rock2^iEKO^* mutant mice and controls produced 100 % surviving mice after 21 days (Figure 7 A-C). No bleeding was detected by inspection of internal organs (not shown), or in retinas of *Rock1^iEKO^* and *Rock2^iEKO^*labeled with IB4 and TER119 at P6 following TAM injection at P1-3 (Figure 7 D-I). Retinal vascular outgrowth was not affected in *Rock1^iEKO^*, but the *Rock2^iEKO^* mutant mice showed a 12% decrease (11.80 ± 1.5) compared to TAM injected littermate controls (Figure S3 A-F). *Rock1^iEKO^*mutants did not show a cell number decrease per area in the retinal vasculature, while *Rock2^iEKO^* mutants showed a significant decrease in cells per area (Figure 7 J-M).

**Figure 7:**
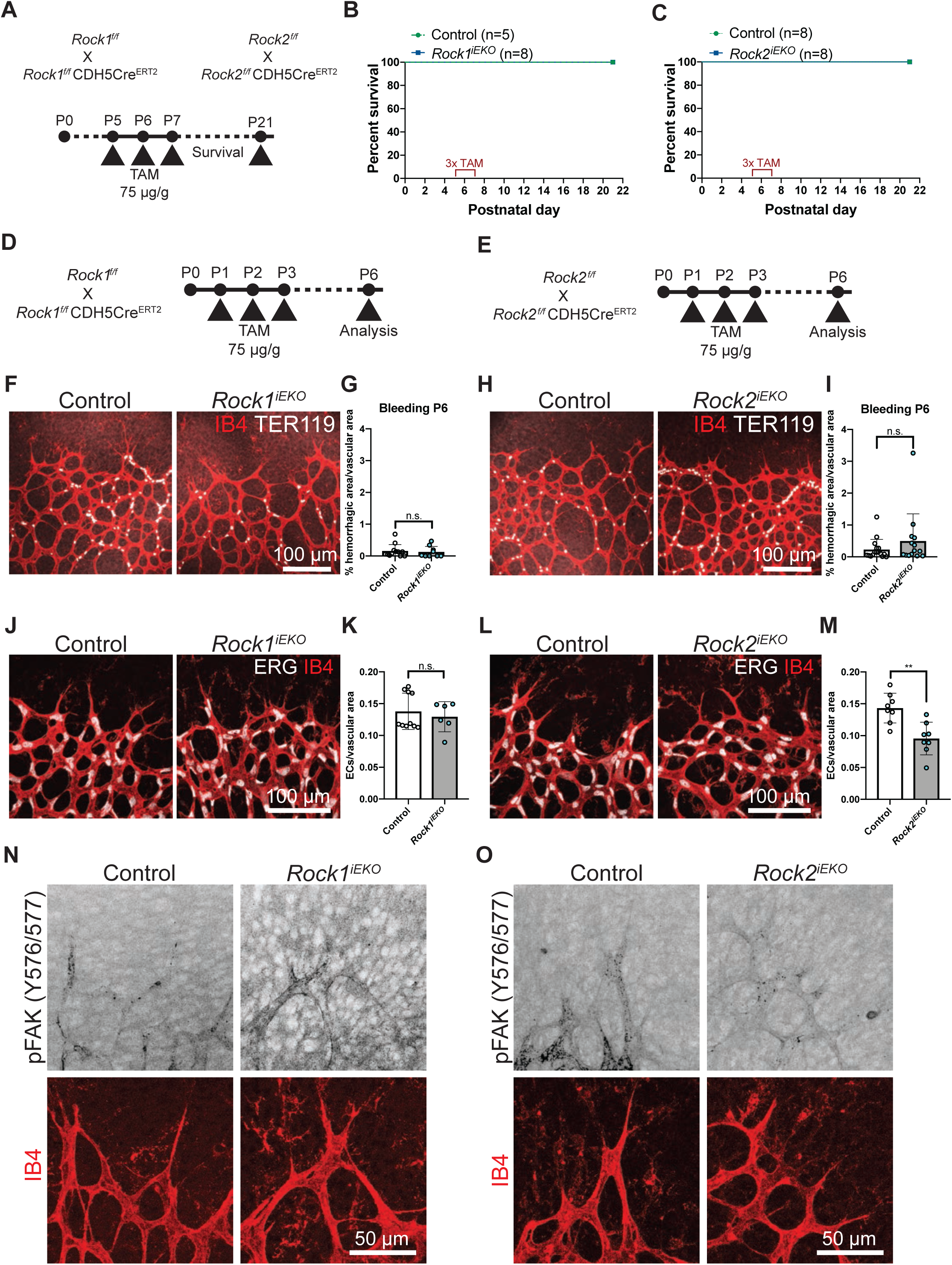
Isoform specific deletion of Rho Kinase in endothelial cells. (A) *Rock1* and *Rock2* gene deletion strategy using TAM injection at P5-7 in postnatal mice. (B-C) Survival curves for *Rock1^iEKO^* (B) and *Rock2^iEKO^* and controls. (C). Green line represents control mice (Cre negative littermates), blue line presents mutant mice. (D-E) *Rock1* and *Rock2* gene deletion strategy using TAM injection at P1-3 in postnatal mice. (F and H) Whole mount retinas from control and *Rock1^iEKO^* mice (F) or control and *Rock2^iEKO^* mice (H) at P6, stained with IB4 and TER119. (G and I) Quantification of bleedings presented in % of hemorrhagic area normalized to vascular area, at P6 for control and *Rock1^iEKO^* (G) or control and *Rock2^iEKO^*(I). (J and L) Whole mount retinas from control and *Rock1^iEKO^* mice (J) or from control and *Rock2^iEKO^* mice (L) at P6 stained with IB4 and ERG. (K and M) Quantification of endothelial nuclei per IB4+ area, at P6 for *Rock1^iEKO^* (K) or *Rock2^iEKO^* (M). Black lines indicate the mean value s.d.; n.s = not significant; **p<0.01; Groups were compared by Mann-Whitney U test. (N-O) Whole mount retinas from control and *Rock1^iEKO^* mice (N) or control and *Rock2^iEKO^*mice (O) at P6, stained with IB4 and pFAK.

To confirm involvement of ROCK2 specific substrates such as FAK in angiogenesis in vivo, P5 retinas of *Rock1^iEKO^* and *Rock2^iEKO^* were stained with IB4 and pFAK antibody. The pFAK staining in *Rock1^iEKO^* mice show no clear difference compared to control retinas from littermates (Figure 7 N). However, the retinal vasculature of *Rock2^iEKO^* mice showed a clear decreased in pFAK staining compared to retinas from control littermates (Figure 7 O). These data further support the importance of ROCK2 in ECs as well as the substrate specificity of ROCK2 for FAK.

### One allele of *Rock2* is sufficient to maintain angiogenesis and vascular integrity

Given the phenotypic difference between single and double *Rock* mutants, we queried if a single allele of either *Rock1* or *Rock2* would be sufficient to maintain vascular integrity and angiogenesis. To address this question, we crossed *Rock1/2^iEKO^* with either *Rock1^iEKO^* or *Rock2^iEKO^* mutant mice to generate 3 allele mutants (*Rock1^flox/flox^ Rock2^flox/wt^*and *Rock1^flox/wt^ Rock2^flox/flox^*) and injected the mice with TAM at P5-P7 (Figure 8 A). Interestingly, the 3 allele mutants with one intact *Rock2* allele produced 100 % surviving mice after 21 days (Figure 8 B). However, a fraction of mice with one intact *Rock1* allele displayed lethality between P14 – P18 (Figure 8 C). IB4 and TER119 staining of the retinal vasculature at P6 after TAM injection at P1-3 (Figure 8 D-E), showed no difference in extravasated erythrocytes or angiogenic defects in *Rock1^flox/flox^ Rock2^flox/wt^* mice with one intact *Rock2* allele (Figure 8 F-G and Figure S3 G-J). However, *Rock1^flox/wt^ Rock2^flox/flox^*mice present bleedings in the retina at P6 as well as angiogenic defects (Figure 8 H-I and Figure S3 H-L). *Rock1^flox/flox^ Rock2^flox/wt^* mice have normal number of ECs per area (Figure 8 J and K), while *Rock1^flox/wt^ Rock2^flox/flox^* mice present with a significant decrease in cells per area (Figure 8 L and M), which is comparable to *Rock1/2^iEKO^* mice (Figure 4 C and F) and stronger than *Rock2^iEKO^*mutants (Figure 7 L-M). Staining of aorta *en face* preparations after adult gene deletion in 8-week-old mice revealed a step-by-step decrease in the intensity of the phalloidin staining with the weakest staining in *Rock1^flox/wt^ Rock2^flox/flox^*mice compared to control littermates (Figure 8 N and O). Interestingly, only the *Rock1^flox/wt^ Rock2^flox/flox^* mice have disturbed junctions (Figure 8 N), revealing that one intact allele of *Rock2* is sufficient to maintain angiogenesis and vascular integrity. This indicates that the amount of polymerized actin correlates with the ability to form and maintain proper barrier function. These data show a strong correlation of cell density in the vascular plexus as well as the amount of F-actin and the severity of the *Rock* mutant phenotypes.

**Figure 8:**
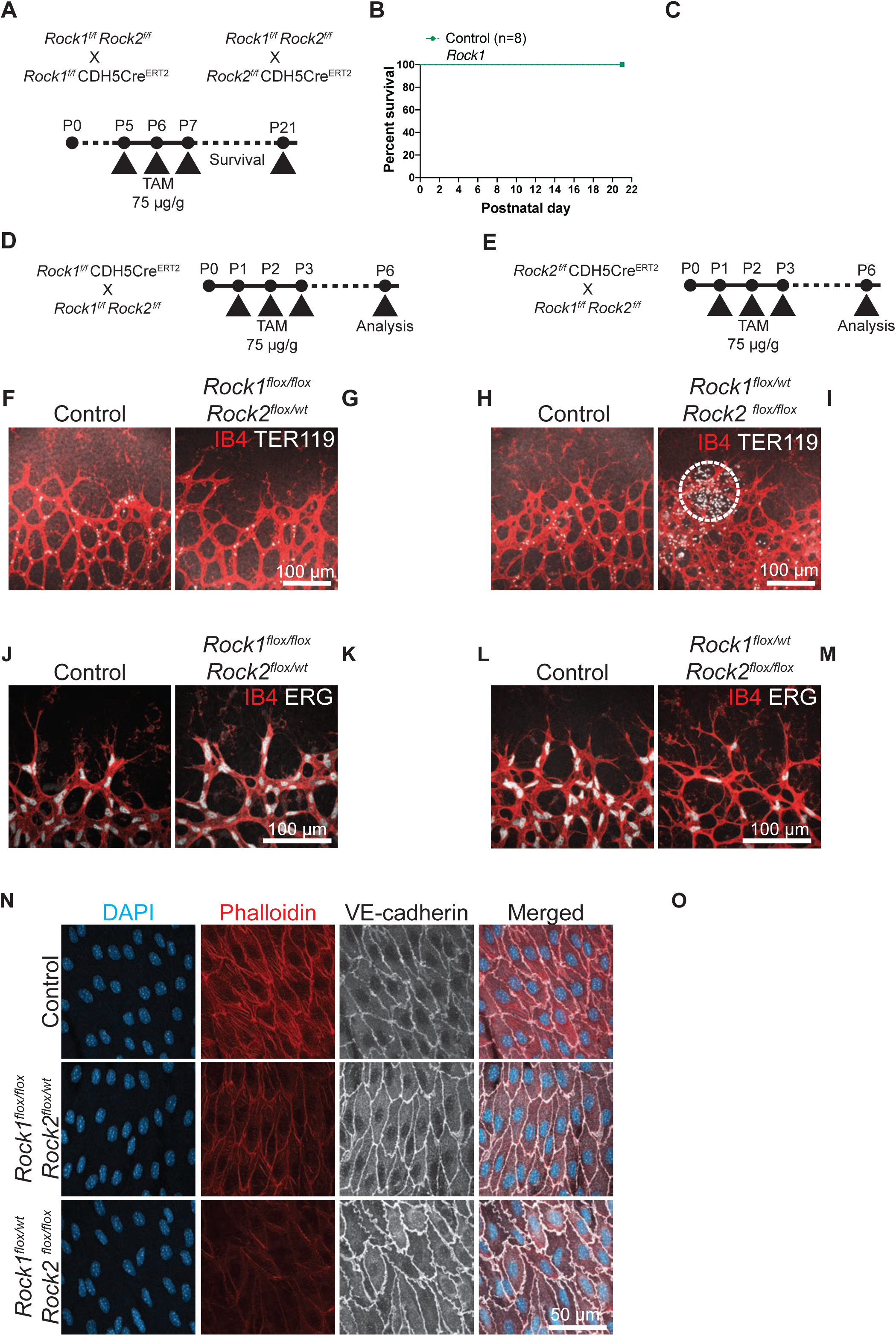
Allele specific deletion of Rho Kinase in endothelial cells. (A) Gene deletion strategy for *Rock*1^flox/flox^ *Rock*2^flox/wt^ and *Rock1^flox/wt^ Rock2^flox/flox^* mice using TAM injection at P5-7 in postnatal mice. (B-C) Survival curve for *Rock1^flox/flox^ Rock2^flox/wt^* (B) and *Rock1^flox/wt^ Rock2^flox/flox^*mice (C); Green line represents control mice (Cre negative littermates); blue line presents mutant mice. (D-E) Gene deletion strategy for *Rock1^flox/flox^ Rock2^flox/wt^* (D) and *Rock1^flox/wt^ Rock2^flox/flox^*mice mice (E) using TAM injection at P1-3 in postnatal mice. (F and H) Whole mount retinas from control and *Rock1^flox/flox^ Rock2^flox/wt^*(F) and *Rock1^flox/wt^ Rock2^flox/flox^* mice (H) at P6; stained with IB4 and TER119. (G and I) Quantification of bleedings presented in % of hemorrhagic area normalized to vascular area, at P6 for control and *Rock1^flox/flox^ Rock2^flox/wt^* mice (G) or control and *Rock1^flox/wt^ Rock2^flox/flox^* mice (H). (J and L) Whole mount retinas from control and *Rock1^flox/flox^ Rock2^flox/wt^* (J) and *Rock1^flox/wt^ Rock2^flox/flox^* (L) mice at P6 stained with IB4 and ERG. (K and M) Quantification of endothelial cells per area, at P6 for *Rock1^flox/flox^ Rock2^flox/wt^* (F) and *Rock1^flox/wt^ Rock2^flox/flox^* (H). Black lines indicate the mean value s.d.; n.s = not significant; ****p < 0.0001; Groups were compared by Mann- Whitney U test. (N) Staining with Phalloidin, VE-cadherin and DAPI of adult mouse aorta after *en face* preparation from control mice, *Rock1^flox/flox^ Rock2^flox/wt^* mice and *Rock1^flox/wt^ Rock2^flox/flox^*mice. (O) Quantification of Actin staining intensity expressed relative units [R.U.]; Dots represent individual pictures; black lines indicate the mean value s.d.; ***p < 0.001; ****p < 0.0001; Groups were compared by ANOVA multiple comparisons test.

## Discussion

This work establishes the crucial role of endothelial ROCK1 and ROCK2 in blood vessel integrity and angiogenesis. We showed that the endothelial specific loss of *Rock1/2* was lethal in postnatal and adult mice, due to multi-organ hemorrhage. Our data indicate that the hemorrhages are a result of decreased F-actin, focal adhesions and disturbed cell-cell junctions. The use of whole mount retina revealed defects in angiogenesis manifested by decreased outgrowth, hyperbranched vascular plexus and decreased endothelial cells per vascular area, resulting in an unperfused vascular front. By using a human sprouting assay, we singled out ROCK2 as the major regulator of these processes in ECs, and that additional loss of ROCK1 further aggravates the observed phenotypes. Furthermore, the sprouting assay revealed changes in actin- and VE-cadherin dynamics resulting in defects in endothelial junction formation and reestablishment as well as cell polarization. Our data indicate that the predominant role of ROCK2 may be due to its substrate specificity towards FAK and LIMK. By carefully analyzing *Rock* single and three allele mutants, we were able to further support the cell culture result and support the importance of ROCK2 in ECs.

This is the first time the role of ROCK1/2 was investigated in ECs with genetic loss of function. Previous studies investigated mostly haploinsufficient mice and did not report early lethality or bleeding [43]. One study generated endothelial specific deletion of *Rock2* using Tie2-Cre mice and conditional *Rock2* “floxed” mice. This investigation showed no impact on body weight, heart rate or systolic blood pressure. However, when these adult *Rock2 -/-* mice were use in an ischemic stroke model, they developed smaller cerebral infarct model compared to controls [44]. The same group investigated *Rock2* deletion in activated fibroblast in the heart using the periostin cre in adult mice. Also, in this model of cell specific *Rock2* deletion in the heart no bleeding or early lethality was reported. Instead, the authors concluded that targeting *Rock2* could protect against left ventricular diastolic dysfunction [45]. *Rock1* was deleted in different neuronal cell types like POMC neurons or AgRP neurons in adult mice and showed no lethality [34,46]. In another study the role of *Rock1* in smooth muscle cells in the context of pulmonary hypertension was investigated using the SM22Cre:*Rock1* KO line. *Rock1* deleted mice still showed no lethality but showed reduced pulmonary hypertension [47]. Our results show no lethality in single endothelial *Rock* mutants. However, the deletion of both alleles results in lethality in postnatal and adult mice soon after the deletion.

The use of ROCK inhibitors is a commonly used tool and has been used to investigate the role of ROCK in endothelial cells. The ROCK2 inhibitor H-1152 has been used to investigate the role of ROCK in retinal ECs in P5 pups and resulted in bleedings similar to *Rock1/2i^EKO^* mice in our study [48]. Interestingly, we do not observe bleedings in any single mutant but only in the double mutant. The different outcomes of these experiments might be explained by the inhibition of ROCK in other cell types or off target effects of the inhibitor. Another group compared the ROCK inhibitors Fasudil and Simvastatin in the application of murine models of cerebral cavernous malformations and were able to decrease the severity of hemorrhages [17]. This can be explained by the known fact that endothelial cells with CCM mutations have increased ROCK activity. Overall, this indicates that the right amount of ROCK signaling is required to maintain vascular homeostasis. Our data further indicate that one allele of *Rock2* is enough to maintain this required amount of ROCK signaling in ECs.

Using transcriptomics, Western blot analysis and immunolabeling *in vivo* and *in vitro,* we were able to show that endothelial deletion of *Rock1/2* results in changes of cytoskeleton, focal adhesions, extracellular matrix and cell junctions. In the past, Tie2-Cre mediated FAK deletion has been shown to result in hemorrhages and vascular defects [49]. The extracellular matrix is a complex structure containing a variety of proteins such as Fibronectin, Laminins and Integrins. Fibronectin is crucial for wound healing, embryonic development and blood vessel morphogenesis [50]. A recent study highlighted the importance of Fibronectin polymorphisms, connecting the genetic variants to increased risk of intraventricular hemorrhages in preterm infants [51]. Some laminins have been investigated in endothelial specific deletion models as well. Tie2-Cre mediated deletion of *Laminin α5* under pathological conditions like intracerebral hemorrhage resulted in enlarged injury volume and elevated BBB permeability [52]. Another group deleted *Laminin α4* using the Pdgfb-Cre line, resulting in excessive spouting angiogenesis, comparable to the hyperbranched phenotype observed in *Rock1/2^iEKO^* mice (Figure S2 A-D) [53]. Furthermore, they linked the effects of Laminin α4 to an interaction with integrins [53]. Integrin β1 was shown to control VE-cadherin localization and blood vessel integrity and the loss of *Itg*β*1* in ECs resulted in bleeding and hyper sprouting [48]. In the same publication, the group showed that deletion of the junctional component VE-cadherin resulted in similar phenotypes. They observed that some but not all the observed defects were phenocopied, including bleeding and angiogenic defects [48]. We speculate that under normal physiological conditions, ECs receive signals from the extracellular matrix that get transmitted into the cell via Integrins to activate Rho kinase, resulting in actin polymerization, focal adhesion formation and overall actin dynamics required for junctional integrity and vascular branching morphogenesis. However, how the cytoskeletal changes that we observed contribute or cause the reported phenotypes *in vivo* is not completely understood. An interesting future direction would be the endothelial specific deletion of *LIMK* or *Cofilin*, which could answer this question.

One major finding of the present study is that ROCK2 is the more important ROCK in ECs, as highlighted by the fact that one allele of *Rock2* was sufficient to prevent any of the observed phenotypes (Figure 8). A possible explanation for this is the substrate specificity of ROCK2 towards FAK and LIMK (Figure 6 L-O). As ROCK1 and ROCK2 share 92 % identity in their kinase domain, they are believed to share most if not all their substrates [54]. However, previous studies identified that ROCK1 but not ROCK2 phosphorylates Rnd2/RhoE in COS7 cells after stimulation with PDGF [55]. Using coimmunoprecipitation with ROCK1 and ROCK2 antibodies in adipocytes and myoblasts showed that ROCK2 but not ROCK1 interacts with IRS-1, indicating differential roles of ROCK1 and ROCK2 in glucose metabolism [56]. Another interesting study showed that the deletion of ROCK1 in MC4R-expressing neurons leads to obesity, which caused by impaired melanocortin action on food intake. However, the authors did not observe these effects in mice lacking ROCK2 in MC4R-expressing neurons [57]. More examples of specificity towards ROCK1 or ROCK2 can be found upstream of the kinases. This includes inositol phospholipids that interact with the ROCK2 PH domain, indicating that ROCK2 could be regulated by PI-3 kinase [7,54]. Overall, there is the possibility that more substrates specific to ROCK1 or ROCK2 exist and that this might be cell specific, stimulus dependent or based on cellular localization.

## Acknowledgements

We thank the Yale Center for Genome Analysis (YCGA) for RNA sequencing.

## Author Contribution

ML and AE designed the study, ML, CF and JF conducted the experiments. JKL and YBK contributed mice. ML, CF, JF and AE wrote the manuscript, and all authors reviewed the manuscript.

## Supplemental Material

### Footnote

#### Nonstandard Abbreviation and Acronyms

EC: Endothelial cell
ERG: ETS-related gene
FAK: Focal adhesion kinase
HUVECs: Human umbilical vein endothelial cells
IB4: Isolectin B4
LIMK: LIM domain kinase
MLC2: Myosin light chain 2
PAK: p21 activated kinase
ROCK: Rho-associated kinase

## Funding

AE 1R01HLI125811

YBK R01 DK129946,YBK

ML was supported by a fellowship from the DFG (431473248)

---

**Figure S1:**
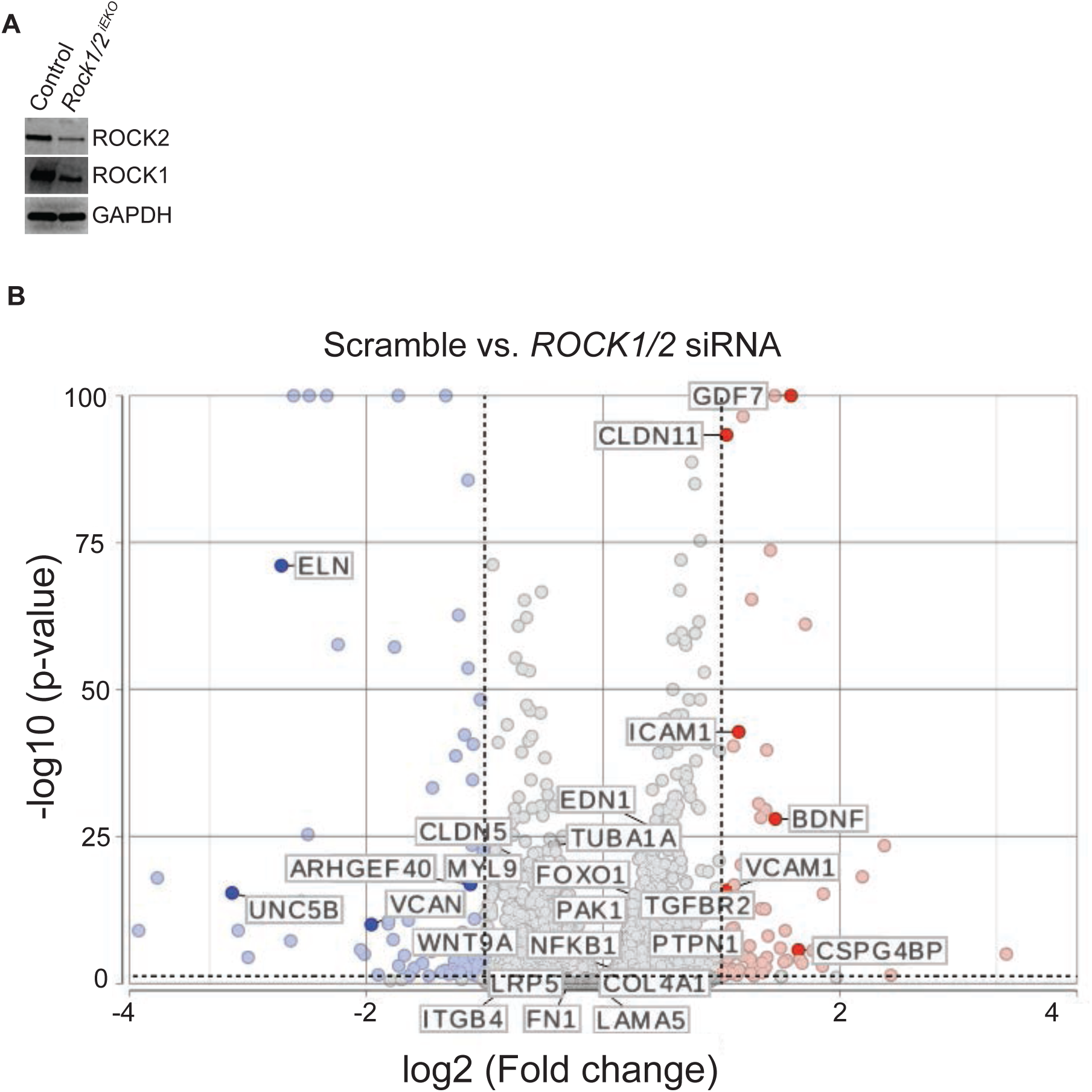
Transcriptional changes after *Rock1/2* knock down in HUVEC. (A) Western blot analysis of protein extract from mouse brain endothelial cell, isolated from *Rock1/2^iEKO^* and Cre negative littermates to validate *Rock* knock out. (B) Volcano plots of differentially expressed genes obtained from pairwise comparison between samples collected from control siRNA treated HUVEC and *ROCK1/2* siRNA treated HUVEC. Volcano plots were generated using log2 fold-change against -log10 (*p*-value) displaying the amount of differentially expressed genes. Genes with 1-fold change or more are highlighted in blue (down-regulated) or red (up-regulated).

**Figure S2:**
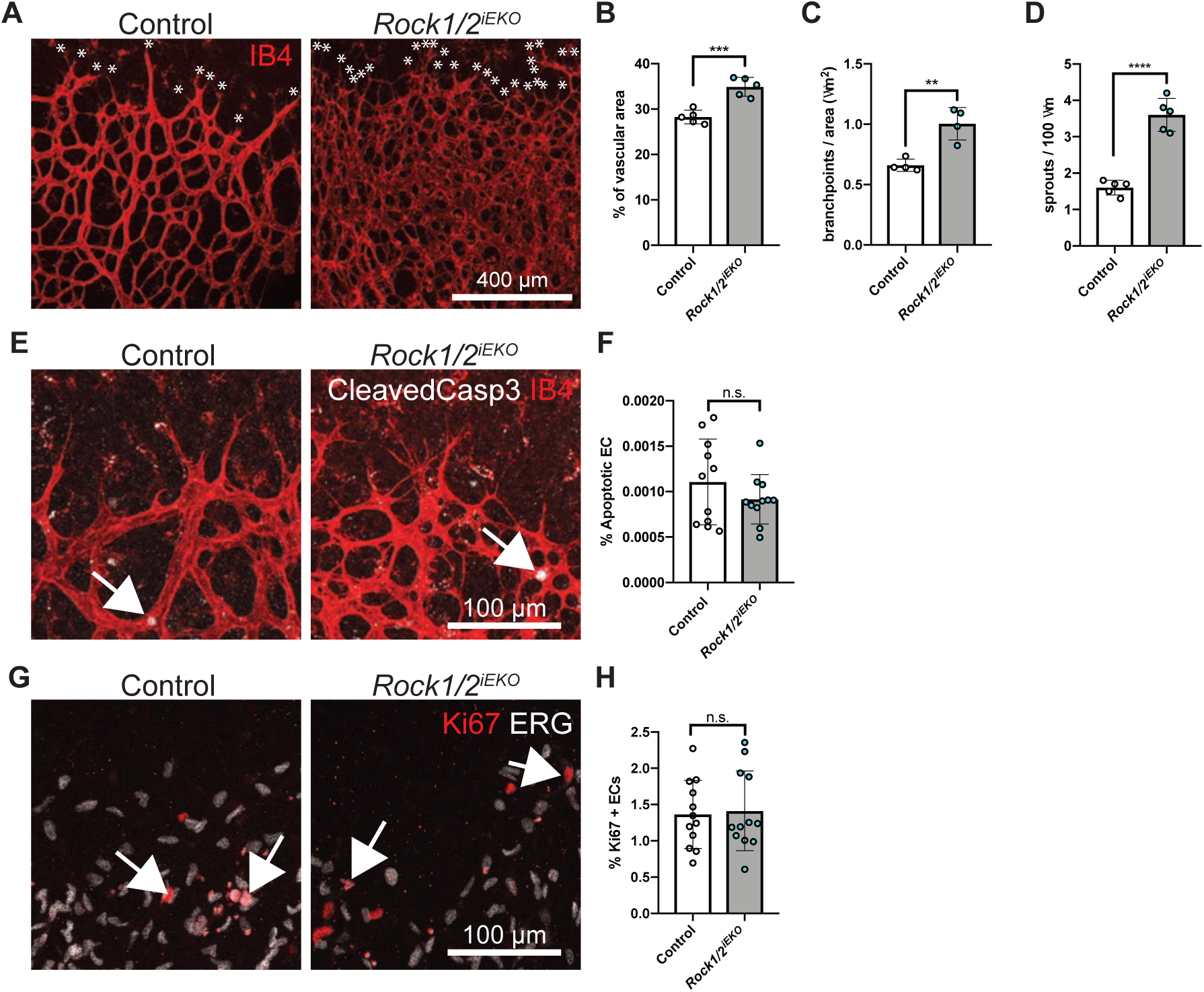
ROCK promotes retinal angiogenesis and blood vessel integrity. (A) Whole mount retinas from control and *Rock1/2^iEKO^* mice at P5 stained with IB4. (B) Quantification of vascular area (B), branchpoints per area (C) and spouts per 100 μm (D) for control and *Rock1/2^iEKO^*. Dots represent individual retinas; black lines indicate the mean value s.d. **p<0.01; ***p < 0.001; ****p < 0.0001. Groups were compared by Mann-Whitney U test. (E) Whole mount retinas from control and *Rock1/2^iEKO^* mice at P5, stained with IB4 and cleaved Caspase3. (F) Quantified % of apoptotic ECs at P5 for control and *Rock* 1/2^iEKO^. Groups were compared by Mann-Whitney U test; black lines indicate the mean value s.d.; n.s = not significant. (G) Whole mount retinas from control and *Rock1/2^iEKO^* mice at P5, stained with ERG and Ki67. (H) Quantification of cell proliferation at P5 for control and *Rock1/2^iEKO^*. Groups were compared by Mann-Whitney U test; black lines indicate the mean value s.d.; n.s = not significant.

**Figure S3:**
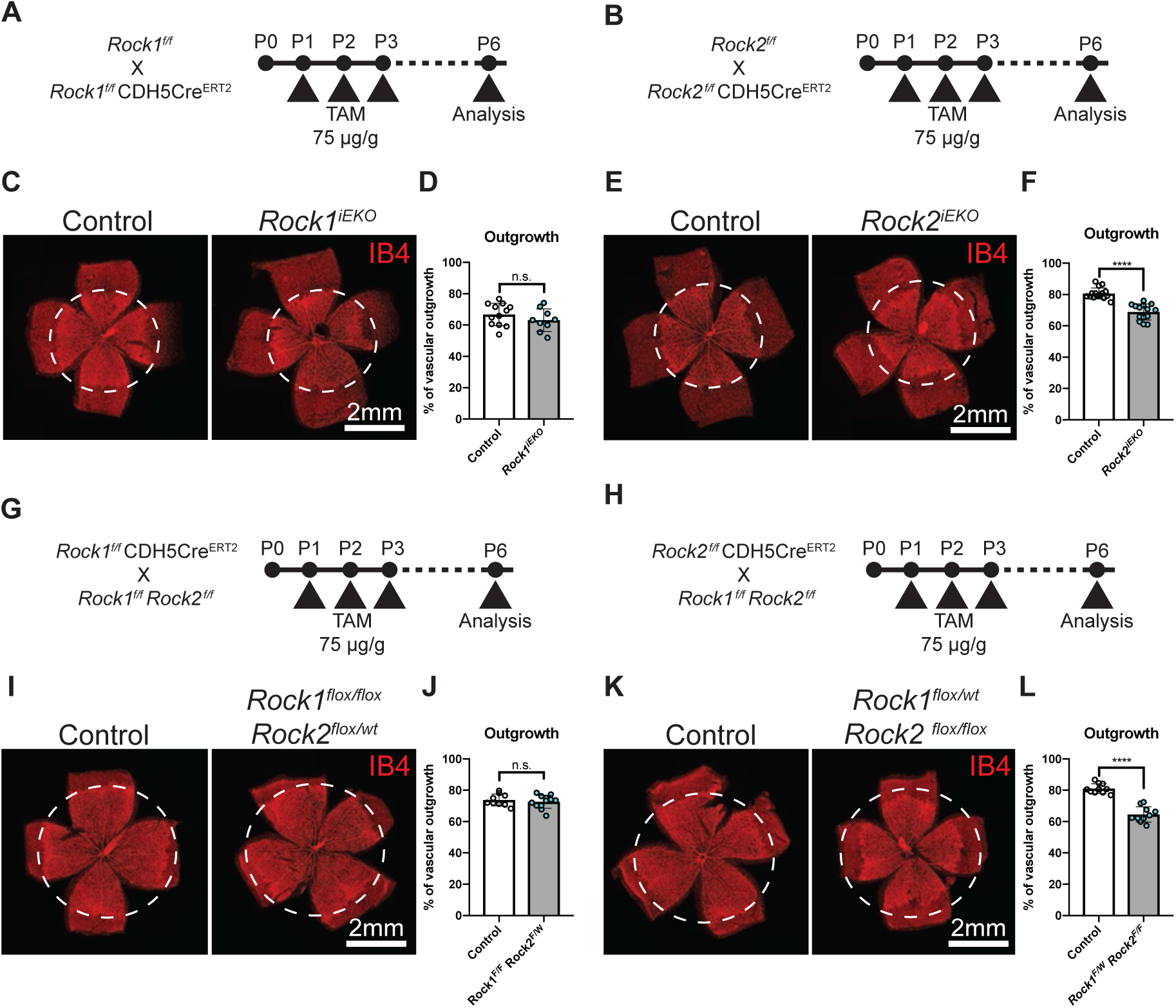
ROCK controls retinal angiogenesis. (A-B) *Rock1* (A) and *Rock*2 (B) gene deletion strategy using TAM injection at P1-3 in postnatal mice. (C and E) Whole mount retinas from control and *Rock1^iEKO^* mice (C) or control and *Rock2^iEKO^*(E) at P6 stained with IB4. (D and F) Quantification of vascular outgrowth at P6 for control and *Rock1^iEKO^* (D) or control and *Rock2^iEKO^*(F). Dots represent individual retinas; black lines indicate the mean value s.d. n.s = not significant; ****p<0.0001. Groups were compared by Mann-Whitney U test. (G-H) *Rock1^flox/flox^ Rock2^flox/wt^* (G) and *Rock1^flox/wt^ Rock2^flox/flox^* (H) gene deletion strategy using TAM injection at P1-3 in postnatal mice. (I and K) Whole mount retinas from control and *Rock1^flox/flox^ Rock2^flox/wt^* (I) or control and *Rock1^flox/wt^ Rock2^flox/flox^* (K) at P6 stained with IB4. (J and L) Quantification of vascular outgrowth at P6 for control and *Rock1^flox/flox^ Rock2^flox/wt^* mice (J) or control and *Rock1^flox/wt^ Rock2^flox/flox^* mice (L). Dots represent individual retinas; black lines indicate the mean value s.d. n.s = not significant; ****p < 0.0001; Groups were compared by Mann-Whitney U test.

